# Epigenetic age acceleration is associated with oligodendrocyte proportions in MSA and control brain tissue

**DOI:** 10.1101/2022.07.20.500795

**Authors:** Megha Murthy, Gemma Shireby, Yasuo Miki, Emmanuelle Viré, Tammaryn Lashley, Thomas T. Warner, Jonathan Mill, Conceição Bettencourt

## Abstract

**Aims:** Epigenetic clocks are widely applied as surrogates for biological age in different tissues and/or diseases, including several neurodegenerative diseases. Despite white matter (WM) changes often being observed in neurodegenerative diseases, no study has investigated epigenetic ageing in white matter.

**Methods:** We analysed the performances of two DNA methylation-based clocks, DNAmClock_Multi_ and DNAmClock_Cortical_, in post-mortem WM tissue from multiple subcortical regions and the cerebellum, and in oligodendrocyte-enriched nuclei. We also examined epigenetic ageing in control and multiple system atrophy (MSA) (WM and mixed WM and grey matter), as MSA is a neurodegenerative disease comprising pronounced WM changes and α-synuclein aggregates in oligodendrocytes.

**Results:** Estimated DNA methylation (DNAm) ages showed strong correlations with chronological ages, even in WM (e.g., DNAmClock_Cortical_, r = [0.80-0.97], p<0.05). However, performances and DNAm age estimates differed between clocks and brain regions. DNAmClock_Multi_ significantly underestimated ages in all cohorts except in the MSA prefrontal cortex mixed tissue, whereas DNAmClock_Cortica_ tended towards age overestimations. Pronounced age overestimations in the oligodendrocyte-enriched cohorts (e.g., oligodendrocyte-enriched nuclei, p=6.1×10^-5^) suggested that this cell-type ages faster. Indeed, significant positive correlations were observed between estimated oligodendrocyte proportions and DNAm age acceleration estimated by DNAmClock_Cortica_ (r>0.31, p<0.05), and similar trends with DNAmClock_Multi_. Although increased age acceleration was observed in MSA compared to controls, no significant differences were observed upon adjustment for possible confounders (e.g., cell-type proportions).

**Conclusions:** Our findings show that oligodendrocyte proportions positively influence epigenetic age acceleration across brain regions and highlight the need to further investigate this in ageing and neurodegeneration.

## 1. Introduction

DNA methylation-based estimators of chronological age, termed as “epigenetic clocks”, make use of penalised regression models such as elastic net regression to select signature CpGs as biomarkers that correlate with chronological age [1, 2]. Multiple clocks have been developed for use as molecular indicators of DNA methylation (DNAm) ages in different human tissues such as peripheral blood [3], multi-tissues (DNAmClock_Multi_) [1], and the brain cortex (DNAmClock_Cortica_) [2]. These epigenetic clocks are being increasingly used to determine the biological age of tissues and organs in various contexts; epigenetic age acceleration has been reported in multiple tissues including the peripheral blood and post-mortem brain tissues in neurodegenerative disorders such as Alzheimer’s disease (AD), Huntington’s disease (HD), and Parkinson’s disease (PD) [4–6].

The DNAmClock_Multi_ has been applied in tissues from several brain regions such as the dorsolateral prefrontal cortex (DLPFC) [6]; frontal, occipital, temporal, and parietal lobes, cerebellum, hippocampus, midbrain, caudate nucleus, and cingulate gyrus [4, 7]; similarly, the DNAmClock_Cortical_ has specifically been tested in dorsolateral prefrontal cortex and posterior cingulate cortex [8, 9]. However, sensitivity of the different clocks to tissue specific changes (e.g., different cell-type proportions) varies depending on the tissues used in the reference training data, with a higher sensitivity likely to be observed in tissue specific clocks compared to pan/multi-tissue clocks [10]. Moreover, when clocks were trained using data from one tissue at a time, their performances in predicting age in another tissue worsened [11]. However, the performance of these clocks in tissues that are particularly enriched for specific brain cell types, such as oligodendrocytes in the case of white matter, remains unexplored.

About half of the human brain is made up of white matter, which primarily comprises myelinated axons and glial cell types such as the astrocytes, microglia, and oligodendrocytes, with oligodendrocytes being by far the most abundant cell type [12]. White matter has been shown to play prominent roles in the normal functioning of the brain including cognitive [13], and motor functions [14]. Many neurodegenerative diseases show white matter changes, including AD, PD, and multiple system atrophy (MSA) [15–17]. MSA is a rare adult-onset fatal neurodegenerative disorder with prominent white matter involvement [15]. MSA belongs to the group of α-synucleinopathies along with PD and Dementia with Lewy bodies (DLB) [18]. However, pathological hallmarks of MSA include the abnormal accumulation of misfolded α-synuclein primarily in the oligodendrocytes rather than in neurons, and these aggregates are termed as glial cytoplasmic inclusions [19]. MSA is a predominantly sporadic disease, with a complex aetiology involving the interplay of genetic, epigenetic, and environmental factors [20]. Recent epigenome-wide association studies have revealed aberrant DNA methylation changes in the MSA post-mortem brain tissue [21–23].

We therefore investigated the performances of DNAmClock_Multi_ and DNAmClock_Cortical_ in white matter from multiple brain regions and compared them with tissues comprising a mixture of grey and white matter in neurologically healthy controls as well as in a disease context. As MSA is a neurodegenerative disease model involving white matter pathological hallmarks, we investigated if patients with MSA exhibited accelerated ageing in the white matter of different brain regions such as the cerebellum, frontal, and occipital lobes, as well as prefrontal cortex grey and white matter mixed tissue.

## 2. Materials and methods

### 2.1 Cohort Description

The first cohort (Cohort 1) consisted of white matter tissue carefully dissected from the cerebellum, and frontal and occipital lobes of post-mortem brain tissues obtained from brains donated to the Queen Square Brain Bank. Cohort 1 comprised a total of 127 samples from MSA cases (N=77) and neurologically healthy controls (N=50), which included 93 samples from a previously published study [21] and 34 newly profiled samples obtained from the same brain bank. Cohort 2 consisted of samples from a publicly available dataset (GSE143157) comprising grey and white matter tissue mix from the prefrontal cortex of MSA cases (N=40) and controls (N=37) [22]. We further included a white matter cohort - Cohort 3, which consisted of control samples (N=9) from the corpus callosum, the biggest white matter structure of the brain, obtained from a publicly available dataset (GSE109381) [24]. As MSA is characterized by oligodendroglial pathology, we included an additional Cohort 4, which consists of sorted SOX10+ (oligodendrocyte-enriched) immunolabeled populations from bulk DLPFC healthy control tissue (N=15) [25]. The demographic details of all cohorts including sample numbers and chronological age are described in Supplementary Table 1.

### 2.2 DNA methylation profiling and data pre-processing

For the newly profiled samples from Cohort 1 (N=34 samples), genomic DNA was extracted from frozen brain tissues using a previously established phenol-chloroform-isoamyl alcohol extraction method. Bisulphite conversion was then carried out with 750 ng of DNA using the EZ DNA Methylation Kit (Zymo Research, Irvine, USA), followed by genome-wide methylation profiling using the Infinium HumanMethylationEPIC Bead Chip (Illumina). Bisulphite conversion and methylation profiling were carried out at UCL genomics, and the raw intensity files (.idat) generated. Methylation profiles of 97 samples (a subset of Cohort 1) generated by Bettencourt et al [21], 77 methylation profiles from the publicly available Cohort 2 [22], and 15 SOX10+ methylation profiles of Cohort 4 [25] had been previously generated using the Infinium

HumanMethylationEPIC BeadChip (Illumina), whereas 9 methylation profiles of Cohort 3 [24], had been previously generated using the Infinium HumanMethylation450K BeadChip (Illumina). All cohorts were subjected to harmonized quality control checks and pre-processing.

Raw intensity files for all cohorts were imported into R using the WateRmelon package [26]. The following preprocessing and quality control checks were performed: (i) raw intensities were visualised to identify and filter out atypical and failed samples, (ii) outlier detection to identify and remove outliers, (iii) bisulphite conversion assessment, where intensities from the probes were converted into percentages and samples with <80% bisulphite conversion were removed, (iv) probe filtering to identify poorly performing probes by assessing the bead counts and detection-values. Samples with >1% probes above the 0.05 detection p-value threshold, probes with a beadcount of <3 in 5% of the samples, and sites for which over 1% of samples showed a detection p-value > 0.05 were discarded. Samples were also discarded if sex predictions did not match with phenotypic sex. Following the pre-processing, dasen normalisation was carried out. Cell-type proportions of neuron-enriched (NeuN+), oligodendrocyte-enriched (SOX10+), and other brain cell type (NeuN-/SOX10-) populations in the bulk tissues were estimated using the CETYGO package (https://github.com/ds420/CETYGO) and a sorted celltype reference dataset as described by Shireby et al. [25]. CETYGO estimates cell proportions and quantifies the CEll TYpe deconvolution GOodness (CETYGO) score of a set of cellular heterogeneity variables derived from a genome-wide DNA methylation profile for an individual sample by capturing the deviation between a sample’s DNAm profile and its expected profile.

### 2.3 DNA methylation age estimation

Normalised beta values were used for the estimation of DNA methylation (DNAm) ages using 2 established epigenetic clocks trained to predict chronologic age, the DNAmClock_Multi_ [1], which employs 353 CpGs curated from analysis of DNA methylation from 51 different tissues and cell-types, and the DNAmClock_Cortica_ [2] designed using 347 CpGs to specifically predict age in the human cortex. DNAm ages for DNAmClock_Multi_ were calculated using the advanced analysis with normalisation, on the online calculator (http://dnamage.genetics.ucla.edu/). The DNAm ages for DNAmClock_Cortical_ were calculated as described by Shireby et al. [2].

### 2.4 Statistics

All statistical analyses were performed in R (ver. 4.1.1). For both clocks, DNAm age acceleration was calculated as: 1) difference between the predicted DNAm age and chronological age, and 2) residuals obtained by linear regression of DNAm age on chronological age and adjusting for possible confounders such as cell proportions (either neuronal or oligodendroglial), and duplicated individuals with multiple datapoints. Standard linear regression models were applied for Cohorts 2, 3 and 4; however, as Cohort 1 consisted of a subset of tissues for multiple brain regions obtained from the same individuals, mixed effects linear regression models were implemented using the lme4 package (https://github.com/lme4/lme4/)[27], where DNAm age was regressed against chronological age and estimated cell proportions as fixed effects and individual was included as a random effect. Correlations between chronological and epigenetic ages for the different groups were calculated using Pearson’s coefficient. Comparisons across groups/brain regions were performed using Kruskal-Wallis test, and pairwise comparisons between the groups (MSA vs controls) within each brain region were performed using pairwise Wilcoxon test with the Benjamini-Hochberg multiple testing correction. Correlations between epigenetic age acceleration (i.e., residuals obtained by linear regression of DNAm age on chronological age) and cell-type proportions (e.g., estimates of oligodendrocyte proportions) were calculated using Pearson’s coefficient.

## 3. Results

### 3.1 Comparison of the performances of DNAmClock_Multi_ and DNAmClock_Cortical_ in white matter from different brain regions and correlations with chronological age

Using data derived from both neurologically healthy controls and MSA, we investigated the performances of both DNAmClock_Multi_ and DNAmClock_Cortica_ in white matter dissected from the frontal and occipital lobes, cerebellum, as well as from the corpus callosum and sorted oligodendrocyte-enriched nuclei (SOX10+) and compared them with prefrontal cortex tissue constituting a mix of grey and white matter. The composition of prefrontal cortex tissue resembles that of the datasets used to train and test both clocks. As expected, DNA methylation ages estimated with both clocks showed strong correlations with chronological ages in all brain regions, for both white matter and mixed tissues (Table 1). Overall, compared to the DNAmClock_Multi_, the DNAmClock_Cortical_ age estimates showed higher correlations with chronological age and lower errors for all cortical and subcortical regions, regardless of whether samples were from white matter or mixed tissues (Cohorts 1-4). The DNAmClock_Multi_ age estimates, however, showed a stronger correlation with chronological age than the DNAmClock_Cortical_ for cerebellar white matter (Cohort 1).

**Table 1:**
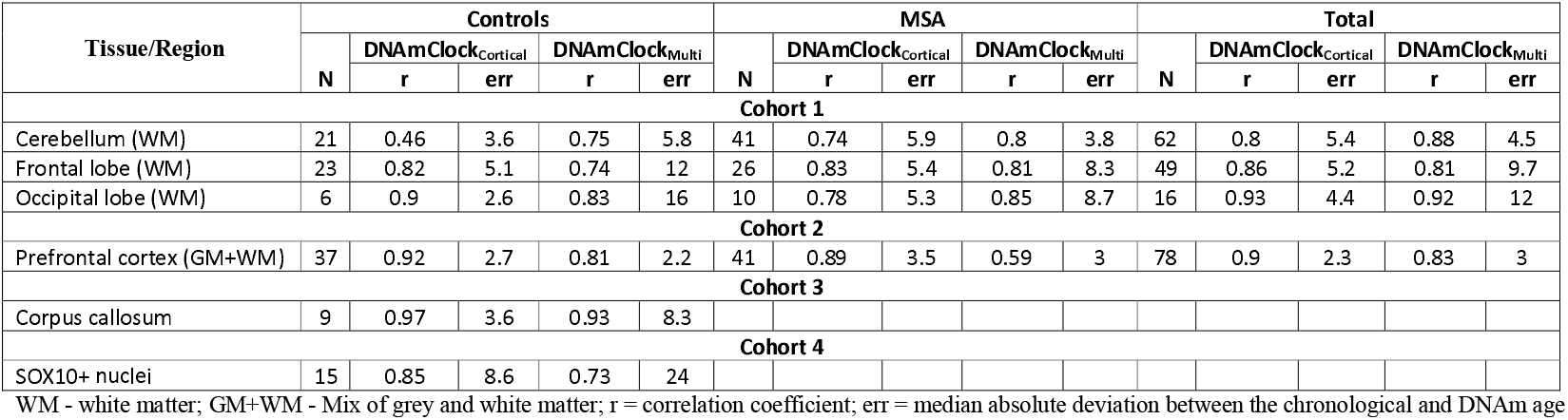
Comparison of correlations between estimated DNA methylation age with DNAmClock_Multi_ and DNAmClock_Cortical_ clocks and chronological age for the different brain regions and cohorts.

For the healthy controls (Table 1), strongest correlation was observed for the corpus callosum (Cohort 3) with DNAmClock_Cortical_ (r=0.97), with a considerably lower error (median absolute difference, error=3.6) when compared to that with DNAmClock_Multi_ (r=0.93, error=8.3). This was followed by the prefrontal cortex grey and white matter mix (Cohort 2), where DNAmClock_Cortical_ showed stronger correlation with similar error values (r=0.92; error=2.7) compared to DNAmClock_Multi_ (r=0.81; error =2.2), followed by white matter from the occipital and frontal lobes (Cohort 1) (Table 1).

In case of MSA, correlations between the chronological age and the DNAmClock_Cortical_ age estimates were in general, similar or weaker than those observed for controls, except in case of cerebellar white matter (Table 1). Moreover, error values were higher in all brain regions in MSA samples compared to that of controls, however, these differences did not reach statistical significance. Conversely, the performance of DNAmClock_Multi_ significantly improved in the white matter tissues of MSA cases compared to controls (Supplementary Table 2). However, the latter could be attributable to the lower chronological ages of the MSA cases (range 50-82 years) compared to controls (range 51-99 years), as previous reports suggest that DNAmClock_Multi_ systematically underestimates DNAm age in individuals over ~60 years old, and systematically overestimates it in individuals below ~60 years old [2].

### 3.2. Epigenetic age estimates vary considerably depending on the epigenetic clock used and the cellular composition of the studied tissues

We then compared the difference in the chronological age and the DNAm age estimates for the control and MSA groups predicted by the two clocks for the different cohorts comprising different brain regions and cellular compositions. Overall, DNAmClock_Cortical_ tended to overestimate age, especially for the MSA cases, with significant overestimations in the frontal lobe white matter and oligodendrocyte-enriched nuclei, whereas DNAmClock_Multi_ showed significant age underestimations in the white matter cohorts (Cohorts 1, 3, and 4) (Fig. 1, Supplementary Tables 3 and 4). The mean ages predicted by both DNAmClock_Multi_ and DNAmClock_Cortical_ were closer to the actual chronological age for both control and MSA groups in the prefrontal cortex mixed tissue (Cohort 2) compared to the white matter cohorts. This result is expected, given this mixed tissue cohort is more similar to the tissue types used to train both clocks. As white matter is highly enriched for oligodendrocytes (up to 70%)[28], we included an additional cohort containing oligodendroglial-enriched nuclei from control individuals to infer the effect of this cell type on epigenetic age estimates. For the oligodendroglial-enriched nuclei, DNAmClock_Cortical_ significantly overestimated the age whereas DNAmClock_Multi_ significantly underestimated the age, similar to that in white matter, but with higher error values (Fig. 1, Table 1). The findings from white matter and oligodendroglial-enriched nuclei suggest that the performances of epigenetic clocks in brain tissue are dependent on the cellular compositions of the tissues used, and that oligodendrocyte proportions might play an important role in the performances of epigenetic clocks in brain tissue.

**Fig. 1.**
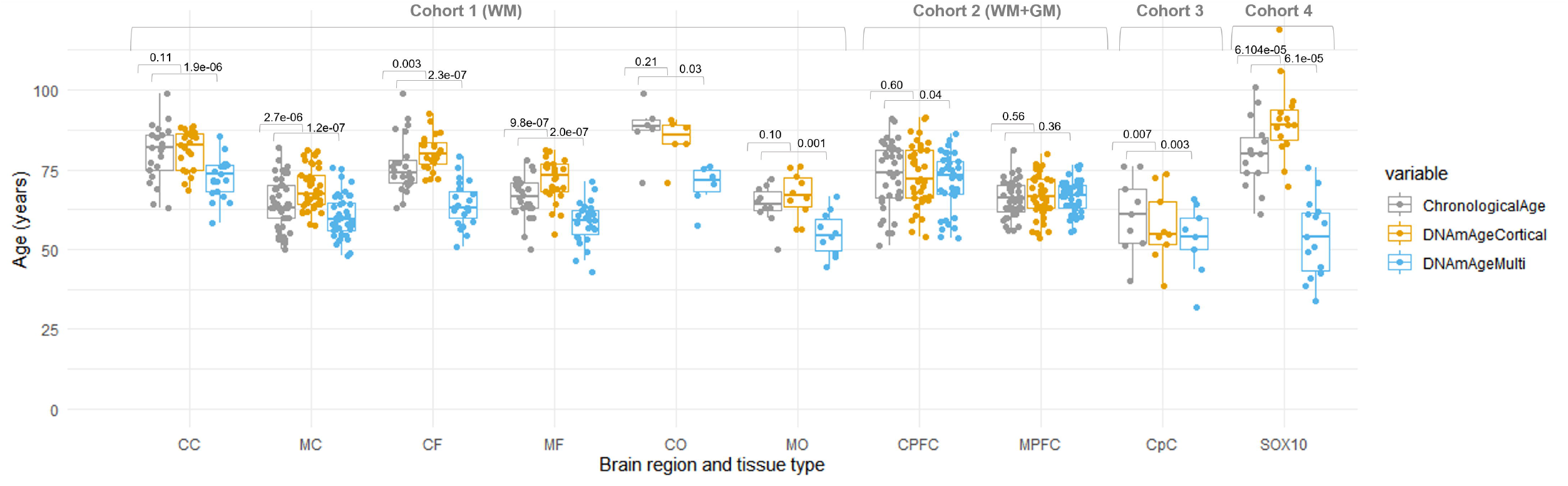
Chronological and DNAm ages for DNAmClock_Multi_ and DNAmClock_Cortical_ for the different brain regions of control and MSA samples in all cohorts. WM – white matter; GM+WM – Mix of grey and white matter; CC – control cerebellum (WM); MC – MSA cerebellum (WM); CF – control frontal lobe (WM); MF – MSA frontal lobe (WM); CO – control occipital lobe (WM); MO – MSA occipital lobe (WM); CPFC – control prefrontal cortex (GM+WM); MPFC – MSA prefrontal cortex (GM+WM); CpC – Corpus callosum; the p-values were calculated using Wilcoxon signed rank exact test for paired samples.

### 3.3 Oligodendrocyte proportions are associated with epigenetic age acceleration

As DNA methylation patterns often are cell-type specific, it is reasonable to hypothesise that differences in specific cell-type proportions in a tissue sample will influence the estimates of epigenetic age based on DNA methylation levels and consequently epigenetic age acceleration. Previous work has demonstrated that significant negative correlation exists between epigenetic age acceleration and the proportion of neurons in the prefrontal cortex [29]. We found the same direction of effect in our prefrontal cortex grey and white matter mix tissues with the DNAmClock_Cortical_, with a significant negative correlation observed between epigenetic age acceleration and neuronal proportions (r=-0.47, p=2.4×10^-10^) (Supplementary Fig. 1). As expected, in our white matter cohorts the estimated neuronal proportions were minimal in frontal (mean 1.8±0.6) and occipital lobes (mean 1.98±1.6) as well as in the corpus callosum (mean 5.7±0.8). Furthermore, there was no relationship between the residual neuronal proportions and epigenetic age acceleration in the white matter for these regions. The opposite was seen in the cerebellar white matter samples, where higher neuronal proportions were found, particularly in the MSA cases (mean 12.8±12.7 vs mean 7.8±11.6 in the controls), and significant negative correlations were observed between the neuronal proportions and the epigenetic age acceleration measures derived from both clocks [DNAmClock_Cortical_ (r=-0.7, p=2.4×10^-10^) and DNAmClock_Multi_ (r=-0.54, p=5.9×10^-6^)] (Supplementary Fig. 1). However, these findings in the cerebellar white matter are not totally unexpected as they may result from: a) the fact that it is challenging to dissect cerebellar white matter without capturing any deep nuclei, particularly in the MSA cases where the cerebellar white matter is very atrophic, increasing the likelihood of deep cerebellar nuclei contamination; and b) the fact that the algorithm used to deconvolute celltype proportions utilizes reference datasets derived from cortical regions and this may influence cell proportion estimates in a region-specific manner.

Lower neuronal proportions (i.e., higher glial proportions) have been associated with higher estimates of epigenetic ages [2] and, in this study, epigenetic ages are largely overestimated in oligodendrocytes (i.e., SOX10+ nuclei) by the DNAmClock_Cortical_. This prompted us to interrogate whether oligodendrocyte proportions would be associated with epigenetic age acceleration. Up to now, proportions of brain cell-types estimates obtained from DNA methylation data were restricted to neuronal vs glial cell-types [with the CETS [30], minfi/wateRmelon packages [26, 31]]. However, the glial group encompasses very heterogeneous cell types, from oligodendrocytes to microglia, astrocytes, and other cells. By using a recently refined cell-type deconvolution algorithm [CETYGO [25]], we dissected the proportions of oligodendrocytes from other glial cells in our cohorts. For estimates derived from both clocks, significant positive correlations were obtained between age acceleration measures and oligodendrocyte proportions for cerebellar, frontal, and occipital white matter as well as for prefrontal mixed tissues (Fig. 2). More details on separate correlations for controls and MSA cases can be found in Supplementary Fig. 3. The strongest correlations with DNAmClock_Cortical_ were found for control cerebellar white matter (r=0.7, p=0.00041), followed by control prefrontal cortex mixed tissue (r=0.65, p=1.3×10^-0^.^5^). For the corpus callosum (Cohort 3), the DNAmClock_Cortical_ estimates, in accordance with the other cohorts, showed a moderate positive correlation with the oligodendrocyte proportions, whereas the DNAmClock_Multi_ showed the opposite, although none of these correlations reached statistical significance in this small cohort. Overall, epigenetic age acceleration estimates derived from DNAmClock_Cortical_ showed stronger correlations with oligodendrocyte proportions compared to that with DNAmClock_Mul_t_i_. This may suggest that the estimates from the DNAmClock_Cortical_ are more influenced by the variation in the oligodendrocyte composition of the tissue than those from the DNAmClock_Multi_. Weaker negative correlations were observed between age acceleration measures and other cell-type proportions (NeuN-/SOX10-) for frontal and occipital white matter as well as for prefrontal mixed tissues and no significant correlations were observed for cerebellar white matter (Supplementary Fig. 3). Altogether, our data supports a role for oligodendrocytes pushing towards older epigenetic age estimates, while it would be the other way around for neurons and the other brain cell types.

**Fig. 2.**
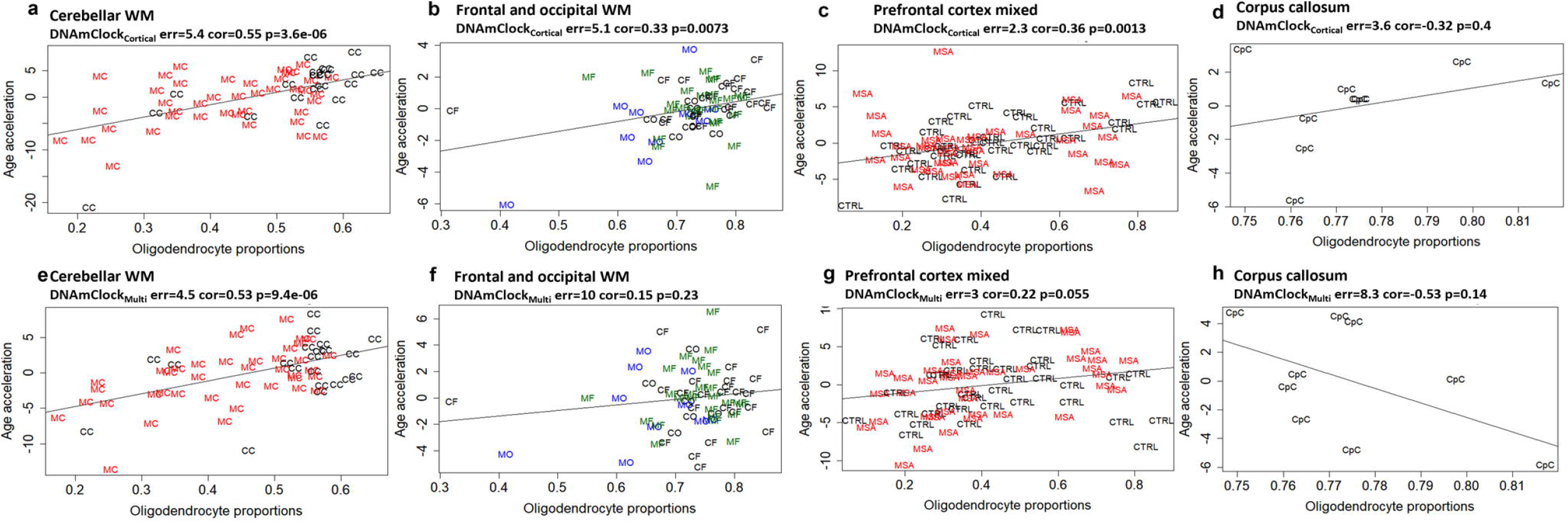
Association between epigenetic age acceleration and oligodendrocyte proportions for DNAmClock_Cortical_ and DNAmClock_Multi_ in the different brain regions. (**a-d**) Age acceleration residuals (y-axis) versus oligodendrocyte (SOX10+) proportions (x-axis) for DNAmClock_Cortica_ in the different brain regions; (**e-h**) Age acceleration residuals (y-axis) versus oligodendrocyte (SOX10+) proportions (x-axis) for DNAmClock_Multi_ in the different brain regions. Age acceleration residuals were obtained by regressing DNA methylation age against confounding factors, including chronological age; oligodendrocyte proportions were obtained using a DNA methylation-based cell-type deconvolution algorithm. The correlation coefficient and p-values shown were calculated using Pearson correlation. CC – control cerebellum (WM); MC – MSA cerebellum (WM); CF – control frontal lobe (WM); MF – MSA frontal lobe (WM); CO – control occipital lobe (WM); MO – MSA occipital lobe (WM); PFC - prefrontal cortex (GM+WM), CpC – Corpus callosum; WM – white matter; GM – grey matter.

### 3.4 Epigenetic age acceleration in the different brain regions in MSA

As increased epigenetic age acceleration has been reported to occur in neurodegenerative diseases [4–6], we investigated the presence of biological age acceleration in MSA when compared to controls based on results from the two clocks. For Cohort 1, with both DNAmClock_Multi_ and DNAmClock_Cortical_, increased age acceleration was observed in the MSA white matter of all brain regions when compared to the controls (Fig. 3). Differences ranged between 2 years in the frontal lobe for DNAmClock_Cortical_ and 7 years in the occipital lobe for both DNAmClock_Multi_ and DNAmClock_Cortical_ (Figure 3A, C). Pairwise comparisons revealed that these differences were statistically significant only in the occipital lobe for DNAmClock_Multi_ (p = 0.042). In the prefrontal cortex mixed tissue (Cohort 2), ~2 years of acceleration was observed in the MSA samples with DNAmClock_Multi_ (p=0.038, Fig. 3B). A smaller difference was found in the same direction with the DNAmClockCortical (Fig. 3D).

**Fig. 3.**
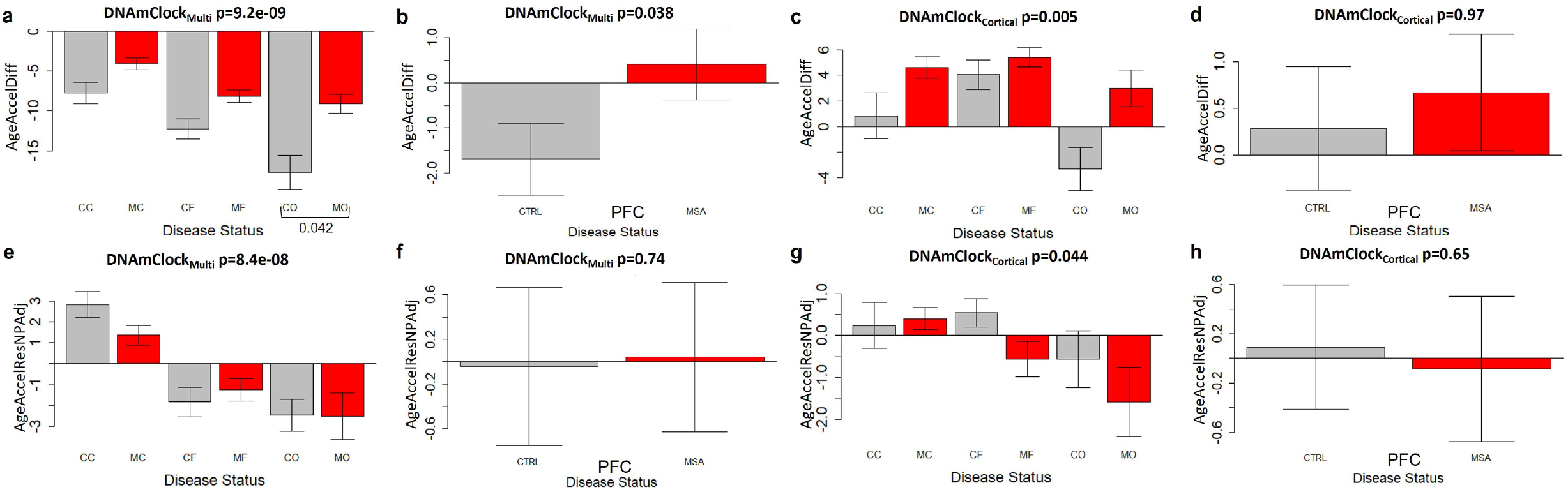
Age acceleration estimates for DNAmClock_Cortical_ and DNAmClock_Multi_ in the different brain regions. (**a-d**)Age acceleration difference for DNAmClock_Multi_ (a - Cohort 1; b - Cohort 2) and DNAmClock_Cortical_ (c - Cohort 1; d - Cohort 2) in the different brain regions, (**e–h**) Age acceleration residual after adjusting for chronological age and neuronal proportions for the DNAmClock_Multi_ (e - Cohort 1; f - Cohort 2) and DNAmClock_Cortica_ (g - Cohort 1; h -Cohort 2) in the different brain regions. CC – control cerebellum (WM); MC – MSA cerebellum (WM); CF – control frontal lobe (WM); MF – MSA frontal lobe (WM); CO – control occipital lobe (WM); MO – MSA occipital lobe (WM); PFC – prefrontal cortex (GM+WM); WM – white matter; GM – grey matter, the p-values for across group comparisons were calculated using the Kruskal-Wallis test and p-values for pairwise analysis between MSA and controls for each brain region were calculated using the Wilcoxon’s test with Benjamini-Hochberg correction for multiple testing.

We then proceeded to verify whether the age acceleration could be influenced by possible confounding factors. Chronological age is known to have an impact on the DNAm age estimations [32]. As chronological ages of the control group were generally higher than that of the MSA group for all brain regions, this had to be accounted for. In addition, for Cohort 1, although white matter tissue was carefully hand-dissected and enriched for oligodendrocytes, neuronal contaminants could still be present, and therefore the neuronal proportions needed to be accounted for. Also, Cohort 1 included multiple brain regions per individual. Therefore, the second measure of age acceleration corresponded to the residuals obtained by regressing DNAm age on chronological age and neuronal proportions as fixed effects and, in addition, by including individuals as a random effect in the case of Cohort 1. Upon adjusting for the abovementioned covariates, no significant age acceleration was observed in the MSA group in any of the brain regions (Fig. 2E-H). Similar results were obtained after adjusting for oligodendrocyte proportions instead of neuronal proportions in these regression models (Supplementary Fig. 4).

## 4. Discussion

This is the first study, to our knowledge, that comprehensively analyses and describes the characteristics of DNAmClock_Multi_ and DNAmClock_Cortical_ in white matter from multiple brain regions. The study compares the performances of the clocks in white matter from subcortical regions, corpus callosum, and cerebellum, as well as in a mix of white and grey matter tissue obtained from the prefrontal cortex and in an oligodendrocyte-enriched population. The study also considered MSA as a disease model involving white matter pathological changes and investigated the presence of increased DNAm age acceleration in different brain regions.

Both epigenetic clocks showed strong correlations with chronological age in both white matter as well as in mixed grey and white matter tissues. Lower error values for the DNAmClock_Cortical_, and stronger correlations in white matter enriched tissues of the corpus callosum, frontal and occipital lobes, and sorted SOX10+ nuclei, as well as for grey and white matter mix tissues of the prefrontal cortex indicate that the cortical clock is highly specific towards the hemispheric cortical and subcortical brain regions in both grey and white matter tissues.

Conversely, stronger correlations with DNAmClock_Multi_ in the cerebellum (white matter) indicates that the multi-tissue clock performs better for regions of the brain other than cortical and subcortical regions. Furthermore, as reported by Shireby et al [2], DNAmClock_Cortical_ shows reduced accuracy when applied to non-cortical regions, which is exemplified by the weaker correlations observed between chronological and DNAm age especially in the cerebellar control samples. Among the cortical and subcortical regions, DNAmClock_Cortical_ outperforms DNAmClock_Multi_, particularly in white matter tissues, with correlation coefficients and error values in ranges similar to those observed previously by Shireby et al. and Grodstein et al. for the dorsolateral prefrontal and posterior cingulate cortices [2, 8].

Lower neuronal proportions (i.e., higher glial proportions) have been previously associated with higher cortical DNAm ages [2]. Recently, studies that identify epigenetic age acceleration in specific cell populations such as sorted neurons and glia have emerged and show that DNAm age of glial cells is significantly higher than that of neurons from the same individual [33]. Our white matter and sorted SOX10+ nuclei cohorts represent oligodendrocyte-enriched populations. The overestimation of epigenetic age by the DNAmClock_Cortical_, coupled with the positive correlations observed between oligodendrocyte proportions and epigenetic age acceleration in both clocks suggests the influence of proportions of this cell-type within tissues on the precision of DNAm age estimators. Furthermore, given that the proportions of other glial cell-types (NeuN-/SOX10-) are negatively correlated with epigenetic age acceleration, further supports the hypothesis that oligodendrocytes are the drivers of the accelerated ageing in glial cells compared to neurons of the same individual previously observed [33]. Therefore, in addition to tissue-specificity, proportions of different cell-types should also be taken into consideration to increase the accuracy of epigenetic clocks. Our results on the varied performances of the two clocks in different tissue types and regions within the brain further support the need for the development of specific clocks for different cell types and tissues as well as clocks specifically tailored in the context of brain ageing and disease as previously suggested [2, 9].

Epigenetic age acceleration in specific brain regions has been reported in other neurodegenerative diseases such as AD and HD [4, 34]; however, no studies exist on epigenetic age acceleration in the brain tissues for any of the α-synucleinopathies. Our rigorous multi-step analysis of the epigenetic clocks in the context of MSA demonstrates that although increased DNAm age acceleration differences were observed in multiple brain regions in the MSA group compared to controls with both epigenetic clocks, no statistical significance was observed upon adjustment for chronological age, neuronal/oligodendrocyte proportions, and duplicated individuals with multiple datapoints. Our results highlight the importance for accounting for confounders during the estimation of age acceleration and emphasize the need for researchers to interpret the results in ageing and disease models with caution.

Our study has several limitations, the sample sizes for each of the brain regions are relatively small, and future experiments with larger sample sizes would be important to further validate our observations; in addition, the chronological age for the controls was considerably higher than that for the MSA cases and although these differences were accounted for during the statistical analysis, different age ranges may have influenced the performance of both clocks differently for MSA cases and controls; finally, both clocks used in this study have not been specifically designed for white matter tissues and follow-up studies with more specialized epigenetic clocks are warranted. Notwithstanding, our data point towards important contributions of oligodendrocytes in brain tissue biological ageing and emphasises the need for additional studies on brain ageing and neurodegeneration focusing on this cell type.

In conclusion, this study demonstrates that both DNAmClock_Cortical_ and DNAmClock_Multi_ showed high correlation between DNAm age and chronological age, even in white matter, expanding the applicability of these clocks. As expected, performances and DNAm age estimates varied considerably between clocks and tissue types. With these available tools, we could not detect changes in epigenetic age acceleration in MSA compared to controls when accounting for known confounding factors. Additional important findings from this study consist of the positive correlations observed between epigenetic age acceleration and oligodendrocyte proportions across brain regions and tissue types, with the opposite being observed for all other brain cell types. This, together with previous reports of older DNAm ages in glial cells compared to neurons from the same individual, support differential biological ageing in different brain cell types, and the possibility of accelerated ageing in oligodendrocytes. Our findings highlight the need for cell type and tissue-specific clocks and clocks that include shared markers of common aberrant epigenomic patterns underlying neurodegeneration to accurately dissect disease related DNAm age acceleration from normal ageing.

## 5. Acknowledgments

The authors would like to thank Dr Sandrine Foti, Ms Gaganjit Kaur Madhan (MSc) for Cohort 1 sample processing, and UCL Genomics centre for advice and processing of the EPIC arrays. Queen Square Brain Bank for Neurological Disorders receives support from the Reta Lila Weston Institute of Neurological Studies and the Medical Research Council. MM is supported by a grant from the Multiple System Atrophy Trust awarded to CB. GS was supported by a PhD studentship from the Alzheimer’s Society awarded to JM. TL is supported by an Alzheimer’s Research UK Senior Fellowship. TTW is supported by the Reta Lila Weston Trust and the MRC (N013255/1). CB is supported by the Multiple System Atrophy Trust and Alzheimer’s Research UK.

## 6. Ethical Approval

Post-mortem brain tissue donated to the Queen Square Brain Bank (QSBB) archives, including the tissue used to generate DNA methylation data for Cohort 1, is stored under a licence from the Human Tissue authority (No. 12198). The QSBB brain donation programme and protocols have received ethical approval for donation and research by the NRES Committee London – Central [18/LO/0721].

## 7. Author Contributions

MM contributed to the experimental work, analysis and interpretation of data, and drafted the manuscript. CB made substantial contributions to the conception and supervision of the work, data interpretation and writing of the manuscript. GS and JM provided data and code for some of the data analysis. GS, YM, EV, TL, TTW, and JM contributed to data interpretation, edited the manuscript and provided critical suggestions. All authors have read and approved the final manuscript.

