## Supplementary for "Epigenetic age acceleration is associated with oligodendrocyte proportions in MSA and control brain tissue"

United Kingdom

### Supplementary Information

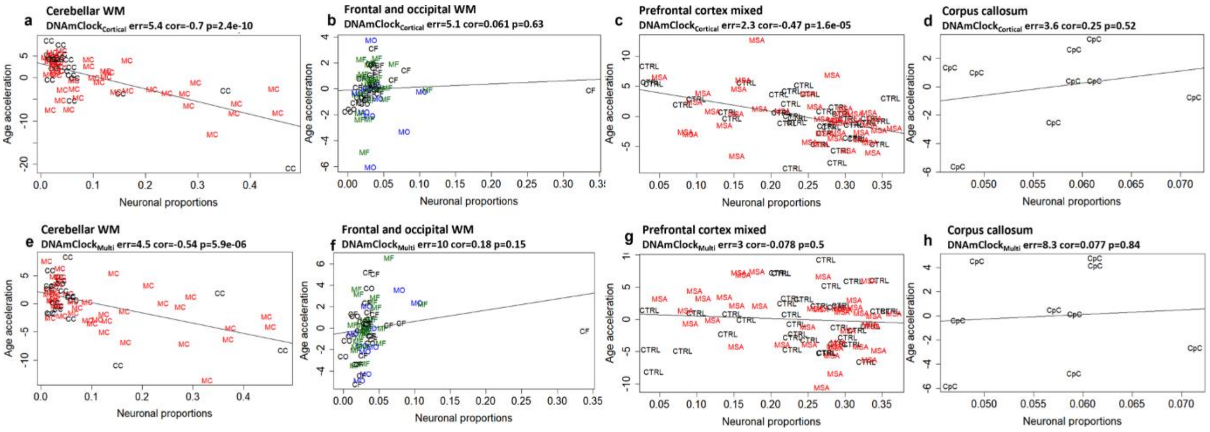


**Supplementary Fig. 1 Association between age acceleration and neuronal proportions for DNAmClock$_{\mathbf{Cortical}}$ and DNAmClock$_{\mathbf{Multi}}$ in the different brain regions. (a-d)** Age acceleration residuals (y-axis) versus neuronal proportions (x-axis) for DNAmClock$_{\text{Cortical}}$ and **(e-h)** Age acceleration residuals (y-axis) versus neuronal proportions (x-axis) for DNAmClock$_{\text{Multi}}$ in the different brain regions. Age acceleration residuals were obtained by regressing DNA methylation age against confounding factors, including chronological age; neuronal proportions were obtained using a DNA methylation-based cell-type deconvolution algorithm. The correlation coefficient and p-values shown were calculated using Pearson correlation. CC – control cerebellum (WM); MC – MSA cerebellum (WM); CF – control frontal lobe (WM); MF – MSA frontal lobe (WM); CO – control occipital lobe (WM); MO – MSA occipital lobe (WM); PFC – prefrontal cortex (GM+WM), CpC – Corpus callosum; WM – white matter; GM – grey matter, the correlation coefficients and p-values were calculated using Pearson correlation.

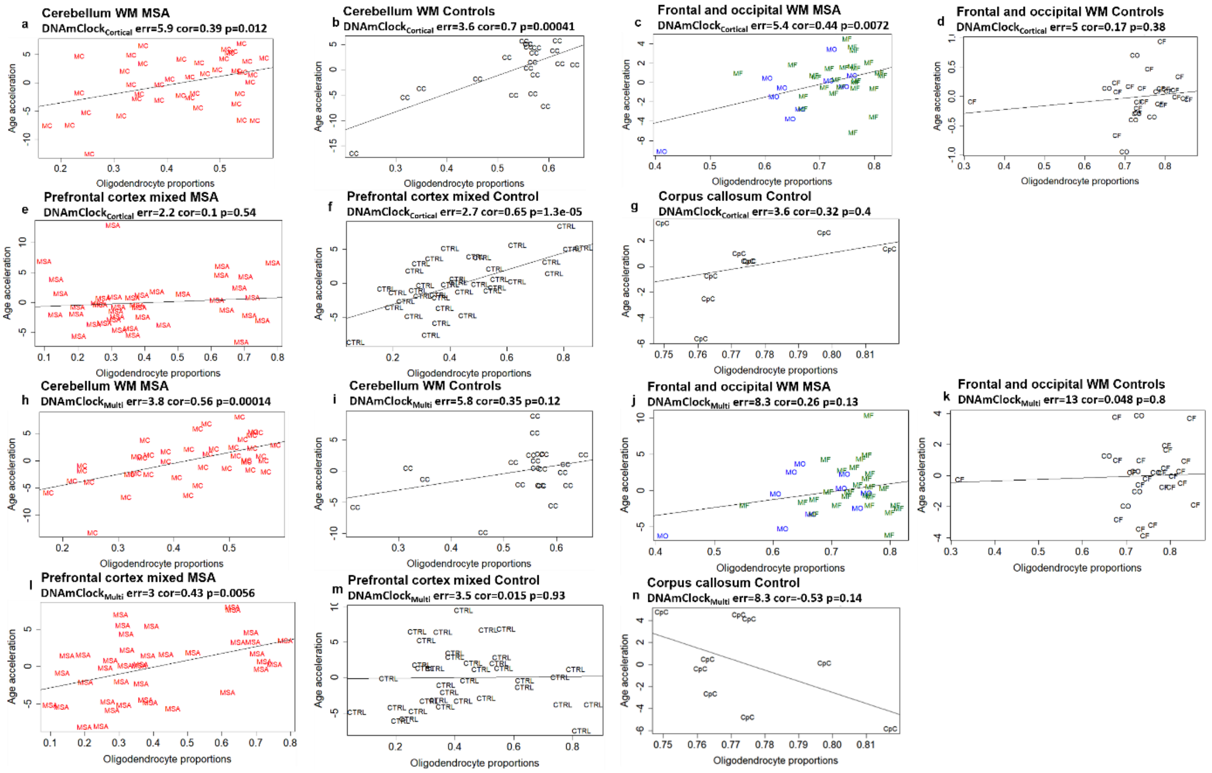


**Supplementary Fig. 2 Association between age acceleration and oligodendrocyte proportions for the MSA cases and control groups for DNAmClock$_{Cortical}$ and DNAmClock$_{Multi}$ in the different brain regions (a-g)** Age acceleration residuals (y-axis) versus oligodendrocyte (SOX10+) proportions (x-axis) for the DNAmClock$_{Cortical}$ and **(h-n)** Age acceleration residuals (y-axis) versus oligodendrocyte (SOX10+) proportions (x-axis) in the different brain regions DNAmClock$_{Multi}$ in the different brain regions. Age acceleration residuals were obtained by regressing DNA methylation age against confounding factors, including chronological age; oligodendrocyte proportions were obtained using a DNA methylation-based cell-type deconvolution algorithm. The correlation coefficient and p-values shown were calculated using Pearson correlation. CC – control cerebellum (WM); MC – MSA cerebellum (WM); CF – control frontal lobe (WM); MF – MSA frontal lobe (WM); CO – control occipital lobe (WM); MO – MSA occipital lobe (WM); PFC - prefrontal cortex (GM+WM); CpC – Corpus callosum; WM – white matter; GM – grey matter, the p-value was calculated using Pearson correlation.

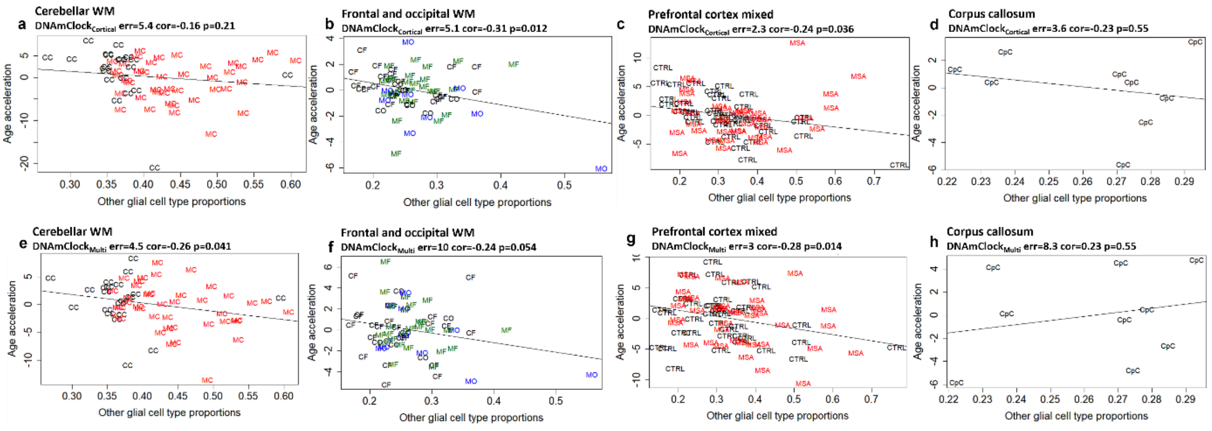


**Supplementary Fig. 3 Association between age acceleration and other glial cell type (NeuN-/SOX10-) populations for DNAmClock$_{Cortical}$ and DNAmClock$_{Multi}$ clocks in the different brain regions. (a-d)** Age acceleration residuals (y-axis) versus other brain cell type (NeuN-/SOX10-) proportions (x-axis) for the DNAmClock$_{Cortical}$ and **(e-h)** Age acceleration residuals (y-axis) versus other brain cell type (NeuN-/SOX10-) proportions (x-axis) DNAmClock$_{Multi}$ in the different brain regions. Age acceleration residuals were obtained by regressing DNA methylation age against confounding factors, including chronological age; NeuN-/SOX10- proportions were obtained using a DNA methylation-based cell-type deconvolution algorithm. The correlation coefficient and p-values shown were calculated using Pearson correlation. CC – control cerebellum (WM); MC – MSA cerebellum (WM); CF – control frontal lobe (WM); MF – MSA frontal lobe (WM); CO – control occipital lobe (WM); MO – MSA occipital lobe (WM); PFC - prefrontal cortex (GM+WM), CpC – Corpus callosum; WM – white matter; GM – grey matter, the correlation coefficients and p-values were calculated using Pearson correlation.

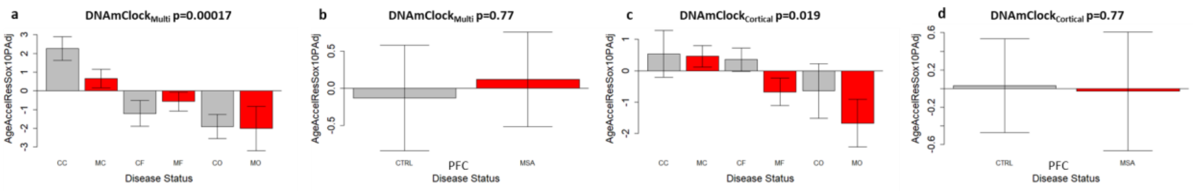


**Supplementary Fig. 4 Age acceleration estimation after adjusting for oligodendrocyte proportions for DNAmClock$_{Cortical}$ and DNAmClock$_{Multi}$ in the different brain regions (a-d)** Acceleration residual after adjusting for chronological age and oligodendrocyte proportions for the DNAmClock$_{Multi}$ (a - dataset 1; b - dataset 2) and DNAmClock$_{Cortical}$ (c - dataset 1; d - dataset 2) in the different brain regions. CC- control cerebellum (WM); MC – MSA cerebellum (WM); CF – control frontal lobe (WM); MF – MSA frontal lobe (WM); CO – control occipital lobe (WM); MO – MSA occipital lobe (WM); PFC - prefrontal cortex (GM+WM); WM – white matter; GM – grey matter, the p-values for across group comparisons were calculated using the Kruskal-Wallis test and p-values for pairwise analysis between MSA and controls for each brain region were calculated using the Wilcoxon's test with Benjamini-Hochberg correction for multiple testing.

**Supplementary Table 1** Cohort demographics of the datasets.

| Tissue/Region | Controls | | | | | MSA | | | | |
|---|---|---|---|---|---|---|---|---|---|---|
| | No. of samples | | | Mean Chronological age ±SD | | No. of samples | | | Mean Chronological age ±SD | |
| **Cohort 1** | N | Females (%) | Males (%) | Females | Males | N | Females (%) | Males (%) | Females | Males |
| Cerebellum (WM) | 21 | 47.6 | 52.4 | 81.1±9 | 79.55±9 | 41 | 48.8 | 51.2 | 64.2±7 | 64.48±8 |
| Frontal lobe (WM) | 23 | 47.8 | 52.2 | 76±10 | 76.58±7 | 26 | 50 | 50 | 65.85±4 | 67.08±8 |
| Occipital lobe (WM) | 6 | 33.3 | 66.7 | 95±5 | 83.75±8 | 10 | 50 | 50 | 64.6±3 | 63.4±8 |
| **Cohort 2** | | | | | | | | | | |
| Prefrontal cortex (GM+WM) | 37 | 48.6 | 51.4 | 72.78±9 | 73.16±11 | 40 | 56.1 | 41.5 | 64.91±5 | 67.47±6 |
| **Cohort 3** | | | | | | | | | | |
| Corpus callosum | 9 | 22.2 | 77.8 | 58±9 | 61.43±13 | - | - | - | - | - |
| **Cohort 4** | | | | | | | | | | |
| Sox10+ nuclei | 15 | 53.3 | 46.7 | 77.88±12 | 81.86±8 | - | - | - | - | - |

WM- white matter; GM+WM - Mix of grey and white matter

**Supplementary Table 2** Difference in median absolute deviation between controls and MSA for $DNAmClock_{Multi}$ and $DNAmClock_{Cortical}$

| | $DNAmClock_{Cortical}$ | | | $DNAmClock_{Multi}$ | | |
|---|---|---|---|---|---|---|
| **Region** | **CTRL** | **MSA** | **p value (two.sided)** | **CTRL** | **MSA** | **p value (two.sided)** |
| **Cohort 1** | | | | | | |
| Cerebellum (WM) | 3.6 | 5.9 | 0.1577 | 5.8 | 3.8 | 0.01412 |
| Frontal lobe (WM) | 5.1 | 5.4 | 0.9123 | 12 | 8.3 | 0.01578 |
| Occipital lobe (WM) | 2.6 | 5.3 | 0.6255 | 16 | 8.7 | 0.007875 |
| **Cohort 2** | | | | | | |
| Prefrontal cortex (GM+WM) | 2.7 | 3.5 | 0.614 | 2.2 | 3 | 0.7949 |
| Total | | | 0.2222 | | | 0.01072 |

WM- white matter; GM+WM - Mix of grey and white matter; the p-values were calculated using two-sided Wilcoxon rank sum test with continuity correction.

**Supplementary Table 3** Mean chronological and DNAm ages for $DNAmClock_{Multi}$ and $DNAmClock_{Cortical}$ for all brain regions of control and MSA samples.

| | N | | Mean chronological age | | Mean DNAm age | | | |
|---|---|---|---|---|---|---|---|---|
| | | | | | $DNAmClock_{Cortical}$ | | $DNAmClock_{Multi}$ | |
| **Tissue/Region** | **CTRL** | **MSA** | **CTRL** | **MSA** | **CTRL** | **MSA** | **CTRL** | **MSA** |
| **Cohort 1** | | | | | | | | |
| Cerebellum (WM) | 21 | 41 | 80.28 | 64.34 | 81.1 | 68.9 | 72.53 | 60.26 |
| Frontal lobe (WM) | 23 | 26 | 76.3 | 66.46 | 80.3 | 71.8 | 64.06 | 58.3 |
| Occipital lobe (WM) | 6 | 10 | 87.5 | 64 | 84.2 | 66.98 | 69.8 | 54.9 |
| **Cohort 2** | | | | | | | | |
| Prefrontal cortex (GM+WM) | 37 | 41 | 72.97 | 66 | 73.26 | 66.66 | 71.27 | 66.41 |
| **Cohort 3** | | | | | | | | |
| Corpus callosum | 9 | | 60.66 | | 57.03 | | 53.17 | |
| **Cohort 4** | | | | | | | | |
| SOX10+ nuclei | 15 | | 79.73 | | 90.11 | | 53.74 | |

WM- white matter; GM+WM - Mix of grey and white matter
